# Plantarflexor fiber length and tendon slack length are the strongest determinates of simulated single-leg heel raise function

**DOI:** 10.1101/415679

**Authors:** Josh R. Baxter, Daniel C. Hast, Michael W. Hast

## Abstract

Achilles tendon ruptures lead to reduced ankle function and often limits recreational activity. Single-leg heel raises are often used clinically to characterize patient function. However, it is unclear how the structure of the Achilles tendon and plantarflexor muscles affects single-leg heel raise function. Therefore, the purpose of this study was to develop a musculoskeletal model in order to simulate the effects of muscle-tendon unit (MTU) parameters on peak plantarflexion during this clinically-relevant task. The ankle joint was plantarflexed by two MTUs that represented the soleus and gastrocnemius muscles. The optimal fiber length, maximal muscle force, muscle pennation, tendon stiffness, and resting ankle angle – a surrogate measure of tendon slack length – were iteratively adjusted to test the combined effects of each of these MTU parameters. Single-leg heel raises were simulated by maximally exciting the two plantarflexor MTUs for each model configuration (N = 161,051 simulations). Optimal muscle fiber and tendon slack lengths had the greatest effect on peak plantarflexion during simulated single-leg heel raises. Simulations that were unable to produce at least 30 degrees of plantarflexion had muscle fibers that were shorter than healthy muscle and longer tendon slack lengths. These findings highlight the importance of preserving muscle fascicle and tendon length following Achilles tendon injuries.

***Funding*** no funding has been provided for this research

***Acknowledgements*** the Authors have no acknowledgements

***Conflict of interest*** the Authors have no conflicts of interest that are relevant to this work

## Introduction

Long-term functional deficits are associated with poor outcomes in many patients following Achilles tendon ruptures.^1^ Improvements in rehabilitation protocols have helped reduced rerupture rates to under 5%, regardless of whether the injury is treated surgically or conservatively.^2^ Despite these improvements in long-term tendon integrity, one-in-five patients do not return to their activities they enjoyed prior to the injury.^3^ Short-term functional deficits are predictive of long-term plantarflexor strength and endurance deficits,^1^ which persist at least seven years following the initial injury.^4^ Despite these documented functional deficits, little is known regarding how muscle-tendon structure dictates function in this patient population. While muscle remodeling^5–7^ and tendon elongation^8^ have been proposed as mechanisms responsible for functional deficits during clinically relevant activities, the relationship between muscle-tendon structure and clinical function are not well understood.

Single-leg heel raise function is a key clinical benchmark for quantifying patient function following Achilles tendon ruptures^9^ and return to activity.^10^ Despite being a simple sub-maximal activity for healthy adults, the single-leg heel raise stresses the plantarflexors of patients recovering from an Achilles tendon rupture. Only one-half of patients are capable of performing a single-leg heel raise 12-weeks after the injury, which is also associated with patient reported outcomes.^9^ Deficits in heel rise height persist at least seven years following tendon rupture^4^ and strongly correlates with tendon elongation over the first year following tendon rupture.^8^ Muscle remodeling following acute Achilles tendon ruptures^5^ and decreased resting ankle plantarflexion^11^ suggest that permanent changes to muscle-tendon unit (MTU) structure governs functional outcomes. Further, this proposed mechanism is supported by similar changes in muscle structure induced by joint immobilization^12^ and changes in tendon travel.^13^

Simple computational models can simulate the effects of small deviations in plantarflexor MTU parameters on ankle and locomotor function. Muscle fascicle length, pennation angle, and peak isometric force are critical parameters that are linked with muscle power.^14–17^ Similarly, tendon stiffness and slack length dictates the shortening demands of the plantarflexor muscles and impacts movement efficiency.^16,18,19^ Resting ankle angle is often measured in clinical settings as surrogate measure of Achilles tendon slack length,^11,20^ which is difficult to measure *in vivo*. While the effects of MTU parameters on walking biomechanics have been studied in great detail,^16,17,21,22^ the multi-factorial implications of clinically-relevant MTU parameters on single-leg heel raise performance is poorly understood.

Therefore, the purpose of this study was to characterize the sensitivity of single-leg heel raise height on plantarflexor MTU parameters. We hypothesized that optimal fiber length and resting ankle angle – a clinical surrogate for tendon length^11,20^ – would have the greatest effect on single-leg heel raise performance. This hypothesis was supported by observations of shorter muscle fascicles,^5^ longer tendons,^8^ and less plantarflexed resting ankle angles^11^ in patients who suffered Achilles tendon ruptures. Because the single-leg heel raise is a sub-maximal activity, we further hypothesized that the potential peak force of the plantarflexors would have less impact on simulation performance. To test this hypothesis, we developed a simple musculoskeletal model that performed single-leg heel raises by maximally exciting the plantarflexor muscles and perturbed several clinically relevant MTU parameters: optimal fiber length, resting ankle angle, pennation angle, peak isometric muscle force, and tendon stiffness.

## Materials and Methods

Simulated single-leg heel raises were completed over a range of MTU parameters of the right plantarflexor muscles (**Figure 1**): optimal muscle fiber lengths, pennation angles, maximum isometric forces, resting ankle angles, and Achilles tendon stiffness values. An open-source musculoskeletal model^23^ (gait10musc18dof) was modified to test the isolated effects of the muscles that crossed the right ankle. The ankle was modeled as a pin-joint that was flexed by a single dorsiflexor muscle, the tibialis anterior, and extended by two plantarflexor muscles, the soleus and gastrocnemius. The soleus muscle is a uniarticular plantarflexor while the gastrocnemius is biarticular, which plantarflexes the ankle as well as flexes the knee. These muscles were modelled as Hill-type muscle bundles comprised of a contractile muscle element in series with an elastic tendon-like element (**Figure 1B**).^24^ Optimal fiber lengths, pennation angles at optimal fiber length, maximal isometric muscle forces, and tendon stiffness parameters were scaled in 10% increments from 50% to 150% of the default model values (**Table 1**). Resting ankle angle varied from 20 degrees to neutral position in 2 degree increments. In total, this parameterization study tested 161,051 combinations of optimal fiber length, pennation angle, maximal isometric force, ankle resting angle, and Achilles tendon stiffness values. Default values for optimal fiber lengths, pennation angles, and maximal isometric muscle forces were adopted from a large cadaveric investigation of architectural properties of lower extremity muscles,^25^ which has been implemented in similar musculoskeletal models.^26^ Tendon stiffness values – which are defined as the amount of tendon strain when loaded with maximal isometric muscle force was set as 4.9%, the default value used in this open-source musculoskeletal model. The ranges in which these MTU parameters were perturbed were based on previous reports of MTU ranges in both healthy and pathologic populations.^5,11,27,28^

**Figure 1 (2column).**
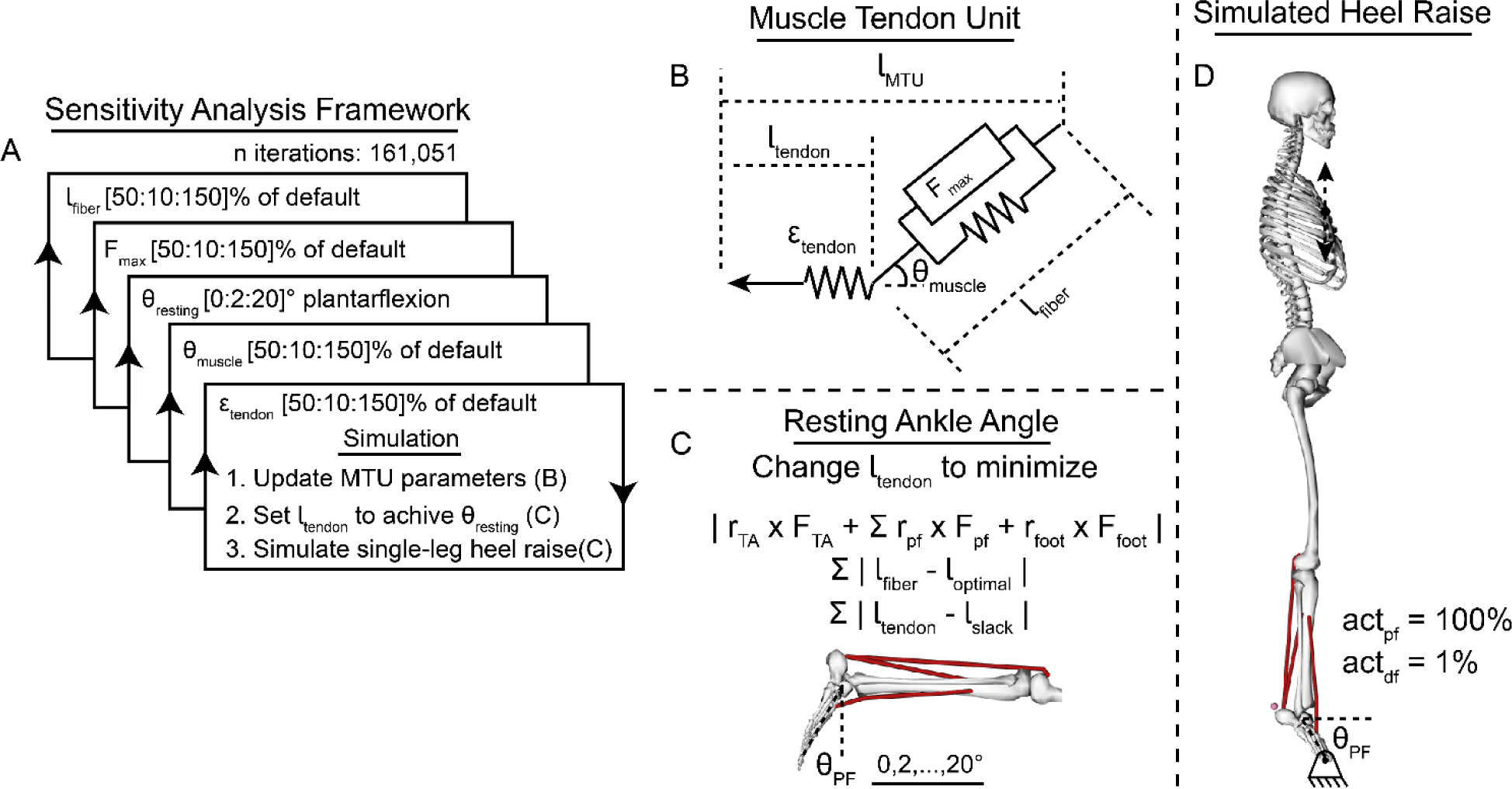
A sensitivity analysis was performed to study the effects of each muscle tendon unit (MTU) parameter on a simulated single-leg heel raise. For each test iteration (A), the MTU parameters were set (B), tendon slack lengths were changed to achieve ankle static equilibrium at the desired resting ankle angle (C), and the single-leg heel raise was simulated (D).

**Table 1.** Default muscle-tendon unit (MTU) parameters

|  | Gastrocnemius | Soleus |
| --- | --- | --- |
| Optimal fiber length (cm) | 5.5 | 4.4 |
| Isometric strength (N) | 1914 | 3586 |
| Pennation angle (°) | 11.0 | 28.2 |
| Tendon Stiffness ($\epsilon$ at isomax) | 0.049 | 0.049 |

Resting ankle angle was achieved by changing the tendon slack lengths of the plantarflexor and dorsiflexion MTUs in order to provide clinical relevance (**Figure 1C**).^11^ We performed the computational analog of instructing the patient to lay prone on a treatment table while the foot and ankle freely hangs at a ‘resting angle’. First, the model was rotated 90 degrees to simulate the subject resting in the prone position, and the optimal muscle fiber lengths, pennation angles, maximal isometric forces, and tendon stiffness values were updated. Second, the ankle angle was set to the desired resting position between 0 to 20 degrees of plantarflexion. Third, the plantarflexor muscle activations were set to 1%, and the dorsiflexor muscle activations were set between 1.5 and 4%. These increased dorsiflexor activation values were selected to balance the plantarflexor muscles, which had a combined maximal force capability of approximately 1.5-4 times that of the dorsiflexor. This was done to achieve ankle equilibrium near the desired resting ankle angle while also maintaining optimal fiber length. Finally, the slack lengths of the plantarflexor and dorsiflexor muscles were iteratively adjusted to minimize a cost function (Eq. 1) comprised of ankle joint torques, differences between optimal fiber lengths and simulated fiber lengths, and differences between the ‘native’ tendon lengths and previously determined tendon slack lengths. Ankle joint torque was calculated as a function of MTU forces and moment arms of both the plantar- and dorsiflexors as well as the weight of the foot segment. The difference between resting muscle-fiber lengths and the prescribed optimal muscle-fiber lengths were minimized to recreate muscle parameters that are measured experimentally using ultrasonography.^29^ This cost function (**Eq. 1**) was evaluated using a gradient-based optimization approach (fsolve, MATLAB, The Mathworks, Natick, MA).

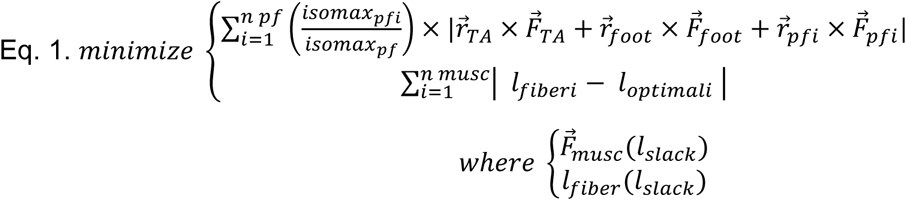

Where l_tendon_ – slack length, *r* – distance from force to ankle joint, *F* – force of MTU or foot segment, *pf* – plantarflexor muscles (gastrocnemius, soleus), *TA* – tibialis anterior, and *musc* – all muscles (gastrocnemius, soleus, tibialis anterior), and F_musc_ and L_fiber_ were both functions of tendon slack length (l_slack_).

Resting ankle angles were confirmed by performing forward simulations to calculate the true resting ankle angle as a function of the musculoskeletal parameters for each parameterized model. To do this, the model was rotated at the pelvis 90 degrees to position the body in the prone position. All joints except for the right ankle were locked to prevent the model from falling and rotating. Plantarflexor activations were held at 1% 3% maximal contraction, while dorsiflexor activation matched the calculated activations for each MTU permutation. Each condition was simulated forward in time for 5 seconds to ensure the ankle came to a complete stop. To confirm that the resting angle was not dependent on the starting position of the simulation, each model configuration was tested at initial positions that were offset from the desired resting ankle angle by 10 degrees plantarflexion and 10 degrees dorsiflexion. These simulations (N = 322,102) always converged on the resting ankle angle (root mean square error: 0.15 degrees).

Single-leg heel-raises were simulated using a simplified musculoskeletal model^23^ to test the implications of MTU parameters on heel-raise height for the 161,051 test permutations (**Figure 1**). Motion constraints were applied to the model to eliminate the need for a control algorithm. The right metatarsophalangeal joint was constrained with the ground using two point constraints in order to limit foot motion with the ground along a foot-fixed medial-lateral axis. The sternum was constrained to move along a vertical axis as a means to ensure the model was in an upright posture during the simulated heel-raise. The joints of the left leg, lumbar, and right knee and hip were ‘locked’ but allowed to move in small amounts (< 0.1 degrees) to satisfy the kinematic constraints of the model. This model constrained knee motion to full extension, effectively converting the gastrocnemius muscle to a uniarticular muscle. Nonetheless, we included both plantarflexors because of the functional differences in MTU parameters^30^ and shortening dynamics during plantarflexion contractions.^31,32^

Forward dynamic simulations were performed to predict the maximal ankle plantarflexion that could be achieved based on the MTU parameters. The gastrocnemius and soleus muscles were maximally excited (100%) while the tibialis anterior muscle was set to 1% of maximal contraction. Each model was simulated for 1 second forward in time. This simulation had a ceiling effect, where some MTU parameters resulted in simulations that over extended the ankle past 75 degrees, which would result in the model jumping. A multivariate linear regression model (fitlm, MATLAB) was utilized to determine the effect a 1% change in each of the MTU parameters. We decided to focus on ankle plantarflexion rather than heel or pelvic vertical displacement to provide greater translation and avoid the effects of stature or foot length. To provide greater clinical relevance to this regression model, we normalized changes to resting ankle angle by the range tested from 0 to 20 degrees plantarflexion. For example, changing the resting ankle angle by 2 degrees was effectively a 10% change in the model.

To further test our hypothesis that tendon length is a primary drivers of single-leg heel raise height, we plotted peak ankle angle as a function of tendon slack length for both plantarflexor muscles and performed univariate linear regression. Since each plantarflexor muscle had unique tendon slack lengths for each simulation, separate linear regressions were performed for each muscle and the tendon slack length was normalized by the range of simulation slack lengths in this study. Tendon slack lengths were normalized by the simulated range for each plantarflexor MTU. The ability to perform a complete single-leg heel raise is a clinical test for patient function following Achilles tendon injuries.^10^ Therefore, we also calculated the implications of MTU parameters on the ability to complete a single-leg heel raise. We set this criteria of a ‘complete’ single-leg heel raise to be 45 degrees of peak ankle plantarflexion, the approximate range of plantarflexion motion under load.^33^

## Results

Peak ankle angle during simulated single-leg heel raises was strongly affected by optimal fiber length (**Table 2, Figure 2**). Despite having no effect on absolute muscle strength, optimal fiber length had a 2.5 times greater effect on heel raise height compared to peak isometric force of the plantarflexor muscles. Additionally, optimal fiber length constrained peak ankle angle, regardless of the other MTU parameters (**Figure 2**). Muscle pennation and tendon stiffness had the smallest effects on peak ankle angle during the simulated heel raise (0.08 and 0.03 degrees for a 1% change in pennation and stiffness, respectively). Eight percent of these simulations fully extended the ankle, which would have resulted in the foot leaving the ground (**Figure 2 top panel**). Similarly, one percent of these simulations were unable to generate any active plantarflexion.

**Table 2.** Effect of 1% change of MTU parameters on ankle angle during simulated single-leg heel raise

| 1% $\Delta$ MTU parameters → | $\Delta$ in peak $\theta_{\text{ankle}}$ |
| --- | --- |
| *Multivariate Linear Regression Model* |  |
| Optimal fiber length | 0.49° |
| Resting ankle angle | 0.31° |
| Peak isometric force | 0.20° |
| Muscle pennation | 0.08° |
| Tendon stiffness | 0.03° |
| *Univariate Linear Regression Model* |  |
| Gastroc. tendon slack length | -0.76° |
| Soleus tendon slack length | -0.81° |

**Figure 2 (2column).**
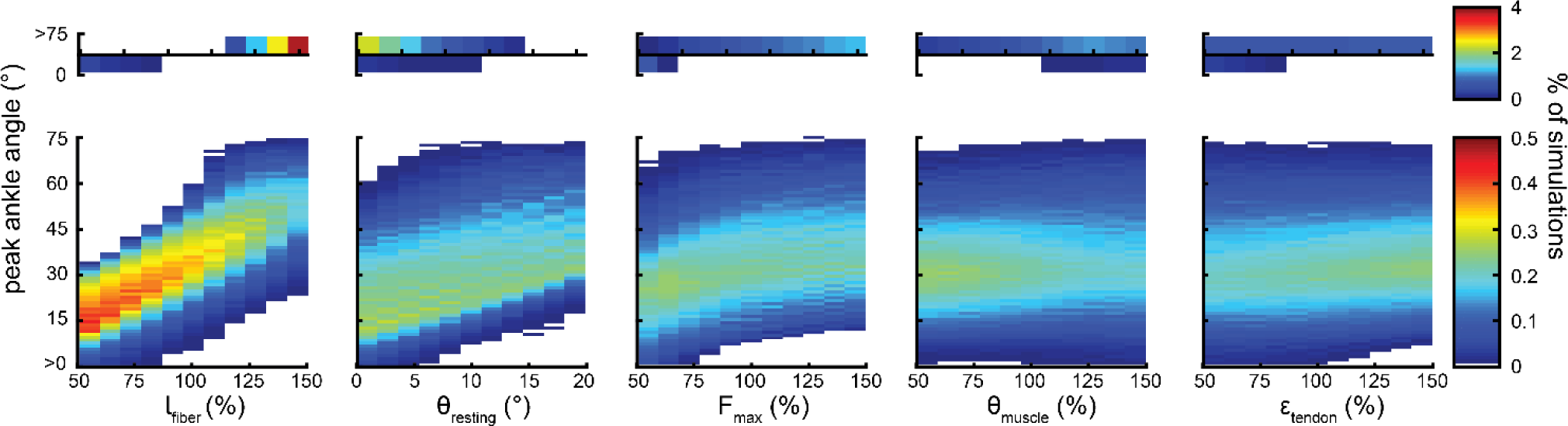
Peak ankle angle during simulated single-leg heel raises were affected by muscle-tendon unit (MTU) parameters (in decreasing order from left to right): optimal fiber length, resting ankle angle, maximum muscle force, muscle pennation, and tendon stifness. Some simulations (*top panel*) resulted in the model plantarflexing past a physiologic range (>75 degrees) or generating no ankle plantarflexion at all (0 degrees).

Tendon slack length is the strongest predictor of peak ankle angle during a single-leg heel raise (**Table 2, Figure 3**). The gastrocnemius and soleus tendon slack lengths ranged from 0.9-1.1 and 0.85-1.13 times their default lengths, respectively. When controlling for these simulated ranges of tendon slack length, a one percent change in the gastrocnemius and soleus tendon slack lengths result in 0.76 and 0.81 degrees decrease in peak ankle angle, respectively (**Table 2**). Gastrocnemius and soleus tendon slack lengths each explained two-thirds of the variability in peak ankle plantarflexion (R^2^ = 0.70 and 0.67, respectively).

**Figure 3 (1column).**
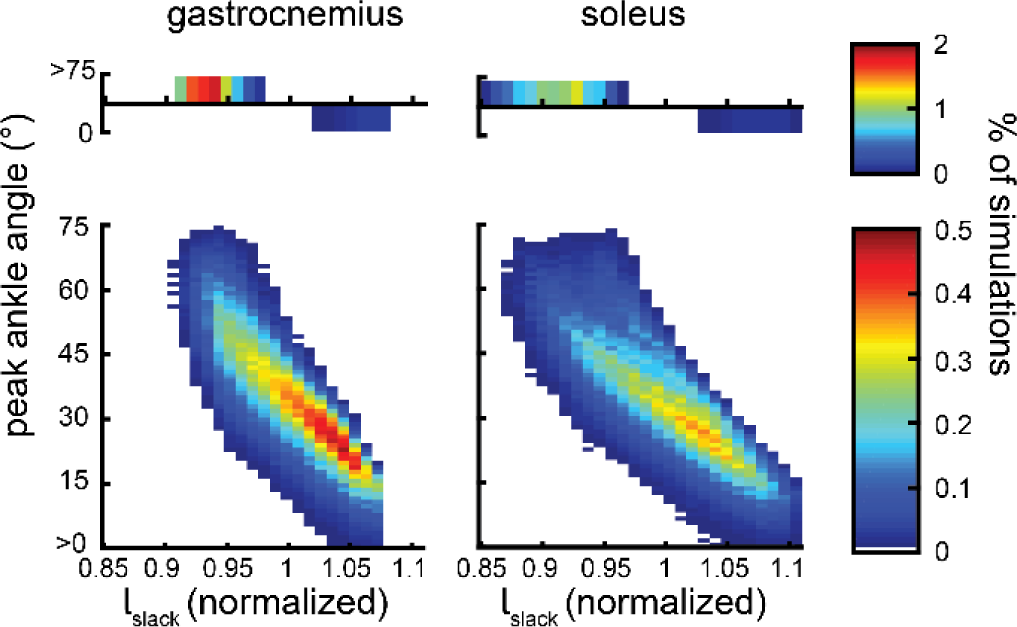
Peak ankle angle during simulated single-leg heel raises were sensitive to small changes in tendon slack length (normalized by default slack length). Shorter slack lengths resulted in greater peak ankle angles while longer slack lengths resulted in smaller ankle plantarflexion *(bottom panel*). Some simulations (*top panel*) resulted in the model plantarflexing past a physiologic range (>75 degrees) or generating no ankle plantarflexion at all (0 degrees).

Two-thirds of the MTU combinations resulted in simulations that did not produce 45 degrees of plantarflexion (**Figure 4**). However, 63 percent of these simulations produced at least 30 degrees of plantarflexion while only eight percent of simulations produced less than 15 degrees of peak plantarflexion. Longer optimal muscle fibers and shorter tendons were most common in simulations that generated at least 30 degrees of peak plantarflexion. Conversely, simulations that were unable to produce at least 15 degrees of plantarflexion had shorter muscle fibers that acted on longer tendons.

**Figure 4 (1.5column).**
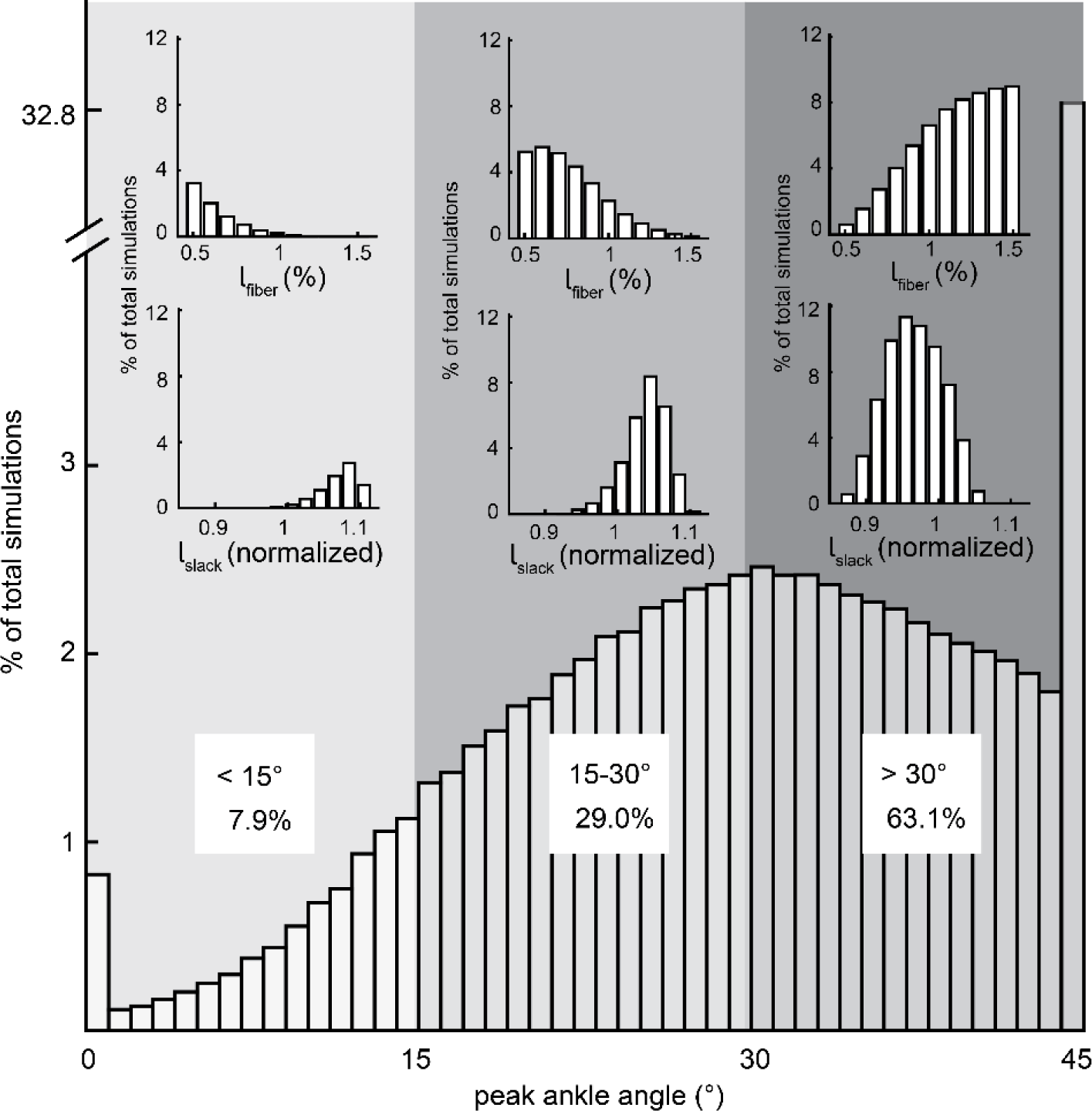
Two-thirds of muscle-tendon unit (MTU) combinations resulted in single-leg heel raises that did not achieve at least 45 degrees of peak ankle plantarflexion. Optimal fiber and tendon slack lengths were the strongest predictors of peak ankle angle. Longer muscle fibers paired with shorter tendons increased single-leg heel raise function.

## Discussion

Achilles tendon ruptures elicit structural and functional changes to the plantarflexor MTU^5,8,27^ that may explain long-term function deficits;^4^ however, the simulated effects of these MTU parameters on clinical function have not yet been defined. Therefore, the purpose of this study was to test the hypothesis that shorter muscle fibers and a less plantarflexed ankle will lead to compromised heel-raise performance. Using a simple musculoskeletal model, we simulated the effects of five parameters that dictate musculo-tendinous function: optimal fiber length, fiber pennation angle, peak isometric force, tendon stifness, and tendon slack length – the latter being calculated using a novel MTU tuning algorithm in order to achieve clinically relevant resting ankle angles.^11^ Our simulation results supported our hypothesis that optimal fiber length and resting ankle angle – the two key factors that dictate the length of the MTU – were the primary determinants of single-leg heel raise function.

The simulation results presented in this study compare favorably with other musculoskeletal simulations of plantarflexion contractions as well as physical measurements of patients with Achilles tendon ruptures. Our findings that optimal fiber length was the strongest predictor of heel raise function agrees with other concentric plantarflexion simulations.^15,34^ The ratio between optimal fiber and tendon slack lengths affected heel raise function (**Figure 3**) similarly to previous simulations of muscle contractions working against a heavy load. Tendon elongation has been measured in clinical populations and explains nearly two-thirds of functional deficits in patients performing single-leg heel raises.^8^ Our simulation findings suggest that tendon elongation coupled with shorter optimal fiber lengths is a likely driver of functional deficits.

Maintaining muscle fiber length while minimizing tendon elongation should be a primary goal when treating acute Achilles tendon ruptures. Limited patient function has been linked to shorter and more pennate plantarflexor muscles^5^ and elongated tendon.^8^ Although surgically repairing the rupture restores resting tendon length, fluoroscopic studies have demonstrated that the rupture site tends to retract and lead to tendon elongation following the first two months of surgery.^35–37^ The magnitude of tendon elongation following surgical repair appears to be sensitivity to early ankle motion following.^35^ Surgical repair also may provide improved long-term plantarflexion power compared to non-surgical treatment of Achilles tendon ruptures.^2^ However, the link between surgical repair and improved patient function is tenuous,^4^ and tendon elongation appears to be a stronger predictor of patient function.^8,38^ Surgically shortening tendon length in a small animal model stimulates muscle remodeling that results in larger, longer, and more powerful skeletal muscle.^39^ Although this prior report highlights the plasticity of skeletal muscle, similar findings in patient populations have not been reported.

Despite the single-leg heel raise being regularly used in the clinic to test patient function following Achilles tendon injuries, this test has a ceiling effect that may limit its descriptive power of patient function. Muscular endurance, measured as the number of consecutive single-leg heel raises performed by a patient has also been used for both clinical research^40^ and guiding return to play decisions.^10^ Tendon elongation has been correlated with deficits in isokinetic ankle power over a range of speeds;^38^ surprisingly, deficits in single-leg heel raise height is more strongly correlated with tendon elongation.^8^ Single-leg heel raises also leverage patient bodyweight as a means to normalize plantarflexor work to the physiologic demands of the patient. Therefore, clinical assessments of patient function following Achilles tendon ruptures should continue to use the single-leg heel raise as a key metric of functional outcomes. Unlike walking and running where energy return through an elastic Achilles tendon is important,^16^ the single-leg heel raise is governed by the amount of positive work done by the plantarflexors. Our simulations suggests that tendon stiffness is not be an important MTU parameter when considering positive work during plantarflexion movements.

This study was affected by several limitations. The musculoskeletal model was a simplified representation of the complicated plantarflexor mechanism. Other factors such as plantarflexor moment arm,^29^ muscle composition,^41^ and foot length^42^ affect plantarflexor function but were excluded in this study to focus on MTU parameters that are documented to change in response to Achilles tendon ruptures.^5,8,27^ The plantarflexors were represented by a single gastrocnemius MTU and a soleus MTU (**Fig. 1**). While other muscles that cross behind the ankle joint have been documented to change their structure following Achilles tendon injuries,^43^ the mechanical advantage and force generating capabilities of these muscles is far less than the primary plantarflexors and were therefore excluded from this study. The multivariate linear regression model calculated the simulated effects of a one percent change in each of the MTU parameters on peak ankle angle during the single-leg heel raise. This calculated effect size is dependent on the range of tested variables. We decided to normalize each MTU parameter by the tested range of each respective parameter. However, each MTU parameter, with the exception of resting ankle angle, were all normalized by the same range and effectively controlled the effect size.

In conclusion, we simulated the effects of plantarflexor MTU parameters on a single-leg heel raise, a common clinical test of patient function. These simulations revealed optimal fiber length and resting ankle angle both had greater effects on peak ankle angle during the heel raise than muscle strength. Treatment plans for acute Achilles tendon ruptures should aim to minimize tendon elongation and muscle fascicle shortening in addition to strengthen the plantarflexors. While surgically repairing the ruptured Achilles tendon allows for tendon length to be modified, the current evidence does not support that this approach provides superior function or outcomes.

